# Transcription of intragenic CpG islands influences spatiotemporal host gene pre-mRNA processing

**DOI:** 10.1101/2020.05.04.076729

**Authors:** Samuele M Amante, Bertille Montibus, Michael Cowley, Nikolaos Barkas, Jessica Setiadi, Heba Saadeh, Joanna Giemza, Stephania Contreras Castillo, Karin Fleischanderl, Reiner Schulz, Rebecca J Oakey

**Affiliations:** Department of Medical and Molecular Genetics, King’s College London, Guy’s Hospital, London, SE1 9RT, UK

## Abstract

Alternative splicing (AS) and alternative polyadenylation (APA) generate diverse transcripts in mammalian genomes during development and differentiation. Epigenetic factors such as trimethylation of histone H3 lysine 36 (H3K36me3) and DNA methylation play a role in generating transcriptome diversity. Intragenic CpG islands (iCGIs) and their corresponding host genes exhibit dynamic epigenetic and gene expression patterns during development and between different tissues. We hypothesise that iCGI-associated H3K36me3, DNA methylation and transcription can influence host gene AS and/or APA. We investigate H3K36me3 and find that this histone mark is not a major regulator of AS or APA in our model system. Genomewide, we identify over 4000 host genes that harbour an iCGI in the mammalian genome, including both previously annotated and novel iCGI/host gene pairs. The transcriptional activity of these iCGIs is tissue- and developmental stage-specific and, for the first time, we demonstrate that the premature termination of host gene transcripts upstream of iCGIs is closely correlated with the level of iCGI transcription in a DNA-methylation independent manner. These studies suggest that iCGI transcription, rather than H3K36me3 or DNA methylation, interfere with host gene transcription and pre-mRNA processing genomewide and contributes to the spatiotemporal diversification of both the transcriptome and proteome.

## INTRODUCTION

Between 20-25,000 protein coding genes have been identified in the human and mouse genomes that give rise to ~200,000 transcripts in tissue- and developmental stage-specific combinations (1). These transcripts can be generated via the use of alternative promoters (2) as well as co-transcriptional pre-mRNA processing mechanisms that include alternative splicing (AS) and alternative polyadenylation (APA) (3–5). Estimates based on transcriptome analyses reveal that ~90% of human transcripts undergo AS (6) and that APA occurs in at least 70% of mammalian pre-mRNAs (7, 8). AS involves the differential inclusion of exons and sometimes introns in the mature mRNA. APA refers to the polyadenylation of transcripts originating from the same gene but that differ in their 3’ end (5). Both AS and APA are dependent on specific sequences recognised by the cellular machinery (9, 10). APA events can occur either at 3’ untranslated regions (UTRs) or intragenic locations. The incidence of both 3’UTR-APA and intragenic polyadenylation (IPA) varies across tissues and cell types providing a way to diversify both the transcriptome and the proteome (11).

Epigenetic modifications of histone tail residues and cytosine bases can influence AS (12–20) and APA (21–23) in developing tissues. Imprinted genes are particularly useful models for the dissection of epigenetic gene expression regulation (22, 23). There are around 130 genes in mouse and 90 genes in human that are subjected to genomic imprinting and are crucial for normal development (24). Monoallelic expression of these genes is coordinated by allele-specific DNA methylation of imprinting control regions (ICRs). Most ICRs acquire differential DNA methylation in the germline (25). Maternally methylated ICRs overlap with promoters, whereas paternal ICRs are found in intergenic regions (25). The active and silent alleles of imprinted genes share the same DNA sequence and are present within the same cellular environment, implying that allelic differences in gene expression are the consequence of epigenetic differences between the alleles (25).

*Mcts2* is an imprinted, monoexonic gene that has resulted from the retrotransposition of *Mcts1* into the fourth intron of the *H13* gene, an event that occurred over 90 million years ago (26). *H13*, referred to as the host gene, is also imprinted. The ICR of *Mcts2/H13* is an intragenic CpG island (iCGI) that overlaps with the promoter of *Mcts2* and undergoes DNA methylation in the female germline (22, 27). As a consequence, *Mcts2* is silent on the maternal allele that generates three *H13* transcripts (*H13a*, *H13b*, *H13c*), which undergo 3’UTR-APA downstream of the iCGI using the canonical polyadenylation sites of the *H13* gene to generate full length transcripts. In contrast, on the paternal allele, the iCGI is unmethylated and transcriptionally active (22, 27). This results in two paternal *H13* transcripts (*H13d*, *H13e*) that undergo intron retention and IPA upstream of the iCGI (Figure 1). The *Mcts2/H13* locus therefore provides a paradigmatic example of how DNA methylation at iCGIs can influence AS and APA.

**Figure 1.**
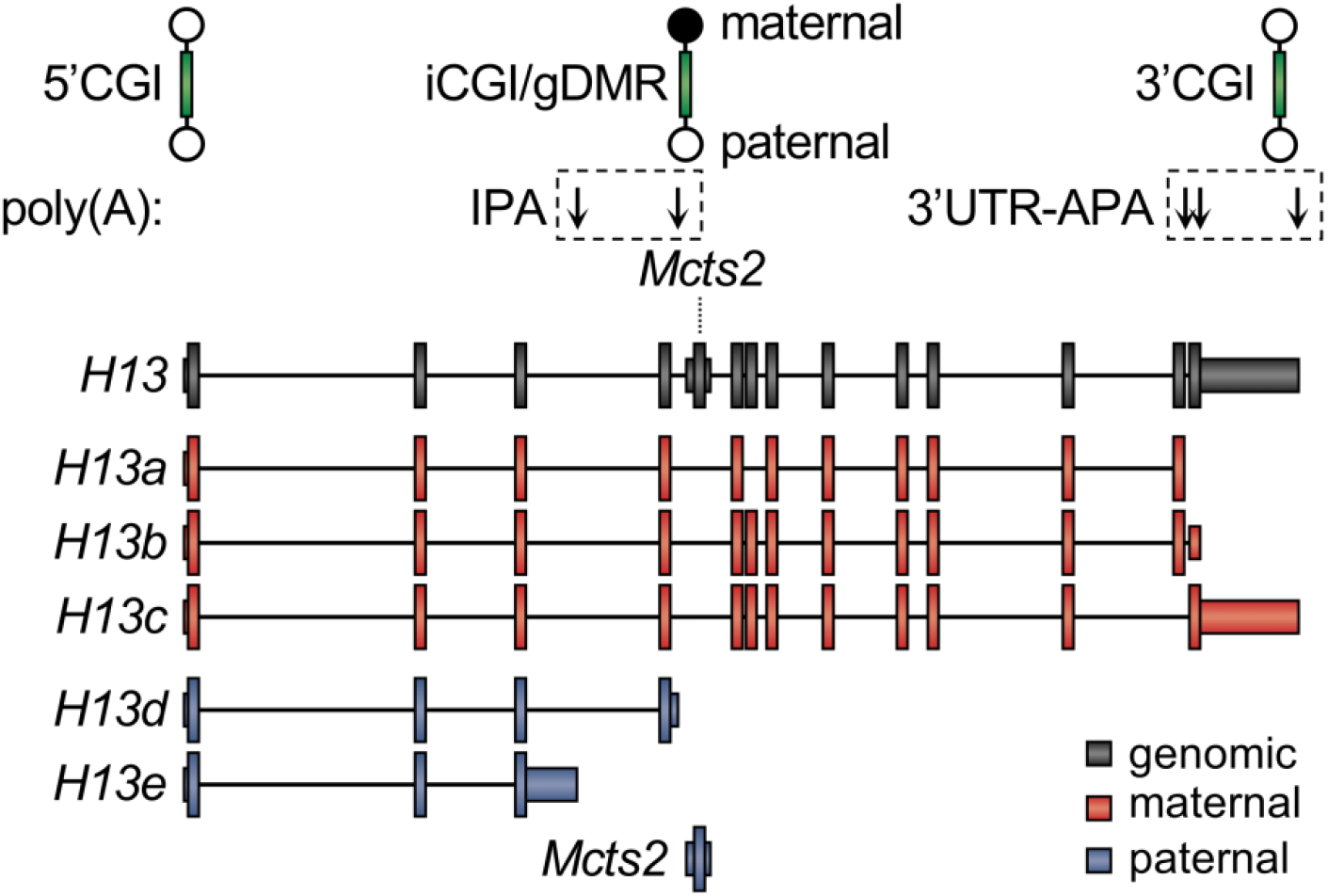
Schematic representation of the *Mcts2/H13* imprinted locus. Three CpG islands (CGIs) (green) are present at this locus: one is associated with the promoter of *H13* (5’CGI), a second with its 3’UTR (3’CGI) and a third, intragenic one with the promoter of *Mcts2* (iCGI). The iCGI is a germline differentially methylated region (gDMR) as it becomes methylated in oocytes. *H13a*, *H13b* and *H13c* are transcribed from the maternal (red) allele and undergo 3’UTR alternative polyadenylation (3’UTR-APA). *H13d*, *H13e* and *Mcts2* are transcribed from the paternal (blue) allele. *H13d* and *H13e* respectively retain portions of intron 4 and 3 and undergo intronic polyadenylation (IPA). Downward pointing arrows represent *H13* alternative polyadenylation sites.

It has been shown that H3K36me3 coordinates tissue-specific usage of alternative exons (14, 19) and prevents intron retention, possibly by facilitating the recognition of weak splice donor sites to ensure introns are correctly spliced out (16, 28, 29). Importantly, H3K36me3 is deposited along the body of actively transcribing genes (30) where it mediates the recruitment of the *de novo* DNA methylation machinery (31) to inhibit spurious transcription initiation from intragenic promoters such as iCGIs (32).

CGIs are regulatory regions typically found at promoters but also at intragenic and intergenic locations. Promoter CGIs (5’CGIs) are generally unmethylated (33) but when methylated, this is usually associated with repression of transcription (34). Biochemical approaches have identified numerous CGIs within host genes (35, 36). iCGIs, which are highly conserved between mouse and human, show tissue-specific patterns of DNA methylation and transcriptional initiation during development, possibly indicating involvement in the spatiotemporal regulation of host gene expression, AS and/or IPA (35–37). The DNA methylation status of these iCGIs is dependent upon their host gene transcriptional activity (33). Host gene transcription across iCGIs is required for the recruitment of *de novo* DNA methylation enzymes and the silencing of said iCGIs. However, the intrinsic capacity of iCGIs to initiate transcription negatively correlates with their sensitivity to this transcription-mediated DNA methylation mechanism (33). The hypermethylated state of iCGIs is associated with low levels of preinitiation RNA Pol II occupancy, H3K36me3 enrichment and lack of H3K4me3 (33). Conversely, iCGIs that retain an unmethylated state show increased preinitiation RNA Pol II binding, are enriched in H3K4me3 and lack H3K36me3 (33). Importantly, when two promoters are located in relatively close proximity, similarly to iCGI/host gene promoters, a transcriptional process initiating from the stronger promoter can have suppressive influence over a second transcriptional process initiating from the weaker promoter, in a phenomenon known as transcriptional interference (TI) (38–41).

Here, we sought to investigate H3K36me3 and transcription as mediators in the generation of alternative transcripts at the *Mcts2/H13* model locus and more broadly. We also sought to determine the influence of intragenic transcription and DNA methylation on host gene pre-mRNA processing. This was achieved through the identification of over 4000 host genes harbouring an iCGI in the mouse genome. The activity of these iCGIs has been found to be tissue- and developmental stage-specific and, for the first time, we demonstrate that the abundance of host gene transcripts terminating upstream of iCGIs is closely correlated with the level of iCGI transcription. These studies suggest that iCGI transcription, rather than H3K36me3 or DNA methylation, interfere with host gene transcription and pre-mRNA processing genomewide, this in turn provides a means to enable spatiotemporal diversification of both the transcriptome and proteome.

## MATERIALS AND METHODS

### Cell culture

NIH/3T3 cells were cultured at 37 °C and 5% CO_2_ in DMEM high glucose (Gibco, 41965039) supplemented with 10% FBS (Gibco, 26140079) and 1X penicillin-streptomycin (Gibco, 15070063). Cells were harvested using 1x trypsin-EDTA (Gibco, 25300054).

### RNA interference

On day one, NIH/3T3 cells were seeded at 15,625 cells/cm^2^ in a well of a 6-well plate. After 24 hours, 3.75 μl Lipofectamine 3000 (Invitrogen, L3000008) was diluted in 125 μl Opti-MEM (Gibco, 31985070). *Setd2*-specific or Scrambled siRNAs (OriGene, SR423523) were diluted in 125 μl Opti-MEM to a final concentration of 10 nM. Diluted Lipofectamine 3000 and siRNAs were mixed and incubated for 20 minutes at room temperature. The siRNA-lipid complex was added to the cells and incubated for 48 hours. Transfections were carried out in triplicate and gene expression was assayed by RT-qPCR.

### Western blot

Protein extracts were obtained by incubating 10_4_ cells/μl in loading buffer (50 mM Tris-Cl pH 6.8, 2% (w/v) SDS, 0.1% (w/v) bromophenol blue, 10% (v/v) glycerol, 100 mM DTT) at 98 °C for 5 minutes. Proteins were separated on 4-12% SDS-PAGE and transferred onto PVDF membrane. Successful transfer of proteins was confirmed by Ponceau S staining. The membranes were blocked in 5%(w/v) TBS-T BSA, incubated with the appropriate primary and secondary antibodies (Supplementary Table S1) and washed with TBS-T. Horseradish peroxidase-conjugated secondary antibody was detected by Pierce ECL Western Blotting Substrate (Thermo Scientific, 32106) coupled with iBright FL1500 Imaging System (Thermo Fisher Scientific, A44241).

### RNA extraction and cDNA synthesis

Total RNA was extracted from pelleted cells using RNeasy Mini Kit (Qiagen, 74104). Membrane-bound genomic DNA was digested with RNase-Free DNase Set (Qiagen, 79254). First strand cDNA was synthesised using a ProtoScript II First Strand cDNA Synthesis Kit (New England Biolabs, E6560) using oligo d(T)23 primers and 500 ng total RNA.

### RT-qPCR

RT-qPCR was performed on a QuantStudio 6 Flex Real-Time PCR System (Applied Biosystems) in a 10 μl reaction including 1 μl cDNA, 1X TaqMan Gene Expression Master Mix (Applied Biosystems, 4369016) and appropriate TaqMan probes (Supplementary Table S2).

#### Statistical analysis

For every condition, three biological and two technical +RT replicates plus one technical-RT replicate were assayed. Relative gene expression was measured using the 2_−ΔCt_ method (42). Statistical significance was determined by unpaired *t*-test assuming consistent scatter and correcting for multiple comparisons using the Holm-Sidak method. Alpha was defined as equal to 0.05.

### mRNA-seq

#### Library preparation and sequencing

After RNA extraction, RNA integrity was measured with 2200 TapeStation (Agilent) using high sensitivity RNA ScreenTape and reagents (Agilent, 5067-5579, 5067-5580 and 5067-5581). TruSeq Stranded mRNA Library Prep (Illumina, 20020594) and TruSeq RNA Single Indexes Set A (Illumina, 20020492) were used to prepare two Scrambled and two *Setd2* knockdown libraries from 900 ng of total RNA per library. Libraries were validated using High Sensitivity D1000 ScreenTape system (Agilent, 5067-5584 and 5067-5585) and KAPA Library Quantification Kit (Kapa Biosystems, 07960140001). All samples were sequenced on one Illumina HiSeq 2500 lane.

#### Differential gene expression analysis

Raw data in the format of FastQ files were subject to quality control using FastQC (0.11.5) (43). Adapter sequences were removed using BBDuk from the BBtools kit (38.22) and the output underwent a second round of quality control by FastQC. mRNA-seq reads were quantified with Kallisto (0.44.0) (44). Differential gene expression (DGE) analysis was carried out using Sleuth (0.30.0) (45). Gene ontology analysis was carried out using PANTHER (14.0) (46) and statistical significance was calculated with the Fisher’s exact test and the Bonferroni correction for multiple testing.

#### Alternative splicing analysis

All mRNA-seq datasets were aligned to the GRCm38 reference genome by HISAT2 (2.1.0) (47) and quality controlled using Picard (2.18.26). MAJIQ (1.1.7a) and Voila (1.1.9) were utilised to detect local splice variants (LSVs) as previously described (48).

### DNA extraction and bisulfite sequencing

Genomic DNA (gDNA) was extracted form pelleted cells using DNeasy Blood and Tissue Kit (Qiagen, 69504). 500 ng of gDNA were bisulfite converted using EZ DNA Methylation-Gold Kit (Zymo Research, D5005).

#### Amplification and cloning

Nested PCR was performed with appropriate primers (Supplementary Table S3) using OneTaq Hot Start DNA Polymerase (New England Biolabs, M0481) and OneTaq Standard Reaction Buffer (New England Biolabs, M0481). Amplicons were cloned into a pGEM-T Easy plasmid (Promega, A1360). Ligation reactions were transformed into chemically competent cells for blue-white colony screening.

#### Sequencing

After colony PCR, amplicons were subject to enzymatic clean-up with ExoSAP-IT PCR Product Cleanup Reagent (Applied Biosystems, 78200.200.UL) and sequenced with T7 and SP6 primers using a BigDye Terminator v3.1 Cycle Sequencing Kit (Applied Biosystems, 4337454) in a 3730xl DNA Analyzer (Applied Biosystems).

#### Methylation analysis

Sanger sequencing results were uploaded to BISMA (49) and the analysis was executed using default parameters.

### Identification of iCGI/host gene pairs

Strand-specific polyadenylated or non-polyadenylated RNA-seq datasets generated by the ENCODE project (50) were utilised. 30 different tissues and/or developmental stages were available for mouse (Figure 5C) and 18 cell lines for human (Figure 5D). Genome assemblies mm9 and hg19 were screened using the Known Genes Canonical table of the UCSC genome browser in conjunction with CGI annotations from Illingworth *et al*. (36). Reads mapping upstream, across and at the iCGI were counted.

### Pearson correlation coefficients

Pearson correlation coefficients (ρ) were calculated using the following formula:

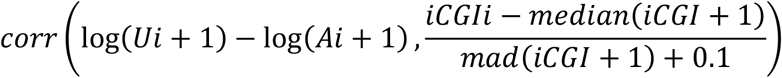

This correlation coefficient measures the linear association between the transcriptional activity of the iCGI and the ratio of host gene expression upstream (U) and across (A) the iCGI calculated across available conditions (*i*). ρ values obtained with this formula were used to generate density plots in Figure 5B and Supplementary Figure S5.

### ChIP-seq data analysis

H3K36me3 and H3K4me3 mouse E14.5 brain ChIP-seq data from the ENCODE project (50) were utilised for average profiling over the host gene body and flanking 3000 bp. Average host gene body length was calculated and divided into three intervals: upstream iCGI; iCGI and downstream iCGI. The logarithm of the fold enrichment over input DNA was calculated at single base resolution for each locus and then scaled to the respective average interval length.

## RESULTS

### Global H3K36me3 depletion does not lead to increased intron retention or intragenic polyadenylation

In mammals, SETD2 is a histone methyltransferase that can interact with the C-terminal domain of phosphorylated RNA Pol II and deposit H3K36me3 along the body of actively transcribed genes with increasingly greater occupancy towards their 3’ ends (51). Previous studies have linked H3K36me3 depletion to increased intron retention in human kidney tumours (28, 29). Intron retention can be used as a proxy for IPA when measured in the context of polyadenylated mRNA since a large fraction of intron retention events map to the 3’ end of transcripts (52). For this reason, only polyadenylated mRNA was selected as a template for the synthesis of cDNA used in gene expression analyses.

To investigate the involvement of H3K36me3 in intron retention and IPA in mouse, NIH/3T3 immortalised mouse fibroblasts were co-transfected with two *Setd2*-specific siRNAs. The efficiency of the knockdown was confirmed by RT-qPCR, which showed an ~80% reduction in *Setd2* mRNA levels in knockdown (KD) samples compared to wild type (WT) samples (Figure 2A). This was further validated by transcriptome analysis (Supplementary Figure S1). As a result, H3K36me3 was globally depleted (Figure 2B). KD cells showed subtle but significant changes in the transcription of 4266 transcripts, of which 1782 (41.77%) were downregulated and 2484 (58.23%) were upregulated (Figure 2C). Among the downregulated transcripts, 432 (10.13%) showed a log2 fold change ≤−1 and 346 (8.11%) of the upregulated transcripts showed a log2 fold change ≥+1 (Supplementary Figure S2A). Gene ontology (GO) analysis using all significantly upregulated genes revealed enrichment of biological processes associated with translation and rRNA metabolism (Figure 2D). Furthermore and consistent with previous findings (53, 54), modification of histones and mRNA metabolism were also among the upregulated biological processes (Supplementary Figure S2B). GO analysis using all significantly downregulated genes in KD samples revealed that the most affected biological processes were DNA replication, cell cycle and cell migration (Figure 2E). SETD2 had previously been shown to methylate α-tubulin (55) and to act as a key regulator of DNA mismatch repair in G1 and early S phase (56). Therefore, increased mitotic and cytokinetic defects were expected. To investigate the role of H3K36me3 in AS, splicing analysis was conducted and resulted in the detection of 136 significant changes in KD samples. However, only three genes (*Ciz1*, *Kctd9* and *Rrm2*) displayed changes in intron retention and in all three cases this type of alternative splicing event was less frequent in the KD than in the WT (Supplementary Figure S3). Additionally, the percentage of intronic bases present in both WT and KD RNA-seq datasets was calculated and the result validated the splicing analysis. Examination of the mRNA-seq metrics from the final quality control step determined the percentage of bases mapping to intronic sequences in WT and KD samples to be 0.111% (± 0.00837) and 0.0994% (± 0.00705), respectively. Taken together, our findings suggest that H3K36me3 does not play a major role in intron retention of polyadenylated mRNA isoforms or IPA in NIH/3T3 cells. These results differ from previous studies that show increased intron retention in human kidney tumours characterised by *SETD2* mutations (28, 29). However, those observations are based on the analysis of RNA-seq libraries generated from total RNA rather than polyadenylated mRNA, which would account for unstable and/or short-lived, intron-retaining transcripts.

**Figure 2.**
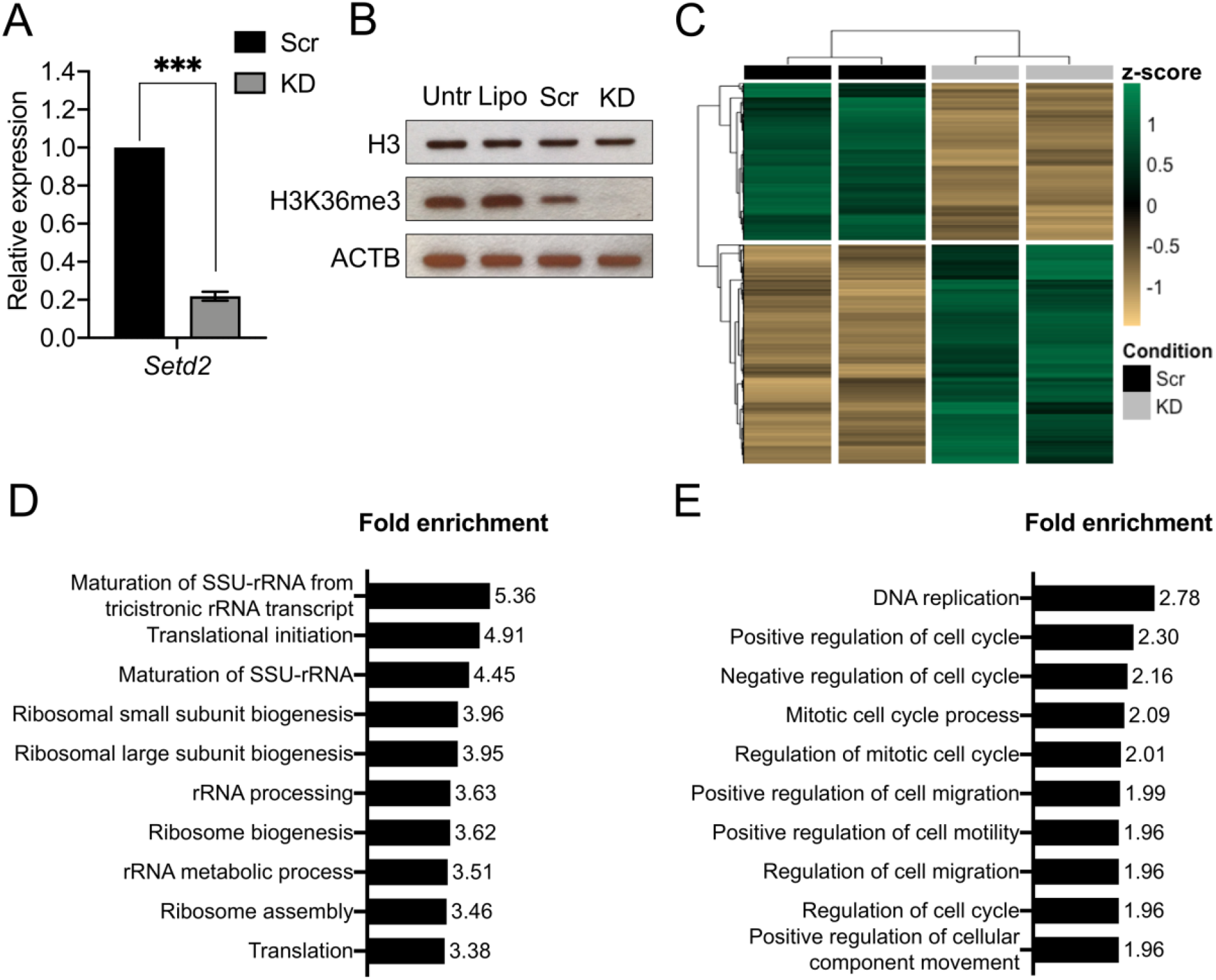
(A) *Setd2* mRNA levels assessed by RT-qPCR 48 hours post transfection. All data are normalised to Ct values for *Actb*. Data are given as mean 2_−ΔΔCt_ values α SEM of three independent experiments. ***, p<0.001 compared with Scrambled group by unpaired *t*-test. Scr, scrambled control; KD, knockdown. (B) Western blot of whole cell lysate showing effective depletion of H3K36me3 upon knockdown of *Setd2*. Total histone H3 levels are unaffected. ACTB was used as a loading control. Untr, untreated cells; Lipo, transfection vehicle only; Scr, scrambled control; KD, knockdown. (C) RNA-seq heatmap of significant differentially expressed transcripts 48 hour post transfection. Values are given as row-wise standard-normalised fragments per kilobase of transcript per million mapped reads (z-score). Scr, scrambled control; KD, knockdown. (D) Top ten upregulated biological processes determined by GO analysis (PANTHER). See Supplementary Table S5 for a complete list of GO ID terms. (E) Top ten downregulated biological processes determined by GO analysis (PANTHER). See Supplementary Table S6 for a complete list of GO ID terms.

More detailed analyses were conducted to investigate a possible locus-specific involvement of H3K36me3 in pre-mRNA processing. Taking advantage of previous studies on the iCGI/host gene pair *Mcts2/H13* (22, 26, 27), this locus was selected as a model system. Using publicly available ChIP-seq data from hybrid immortalised mouse fibroblasts (129/Sv × CAST/Ei) (57), H3K36me3 deposition along the *Mcts2/H13* locus was interrogated. This histone mark is enriched on the maternal (129/Sv) allele at the introns that are retained in the paternally expressed (CAST/Ei) *H13d* and *H13e* transcripts (Figure 3). H3K36me3 has previously been shown to compensate for weak splice donor sites via the recruitment of specific alternative splicing factors (16). When *H13* splice donor sites were scored (58), those present at exon three and four were found to be the weakest compared to the consensus sequence (Supplementary Table S4). It was therefore hypothesised that H3K36me3 may facilitate the recognition and usage of weak splice donor sites at this locus. If this is the case, depleting H3K36me3 would be expected to lead to decreased *H13a, H13b and H13c* isoforms and increased *H13d* and *H13e* isoforms (Figure 1). In other words, depletion of this histone mark would result in an increase in intron retention and IPA. *H13a*, *H13b* and *H13c* (collectively referred to as *H13a-c*) mRNA levels measure 3’UTR-APA, whereas *H13d* and *H13e* expression are a proxy for intron retention and IPA (Figure 1). *H13e* was detectable only at very low levels by RT-qPCR and was not utilised in these expression analyses. Using the same siRNA-treated NIH/3T3 cells as above, RT-qPCR was employed to determine the effects of the knockdown on *Mcts2/H13*. At *H13*, total mRNA levels were reduced by 28% (Figure 4, *H13all*). *H13* isoforms terminating downstream of the iCGI also decreased by 28% (Figure 4, *H13a-c*). This decrease was expected since *H13a-c* constitute the vast majority of *H13* transcripts (22, 27, 59). Contrary to expectations however, *H13d* mRNA levels remained unchanged (Figure 4, *H13d*). Expression from the iCGI was minimally affected (Figure 4, *Mcts2*) and its DNA methylation profile was not altered (Supplementary Figure S4). Taken together, these findings recapitulated those provided by the transcriptome analysis and confirmed that global H3K36me3 depletion via siRNA knockdown approaches may not be sufficient to lead to increased intron retention or IPA.

**Figure 3.**
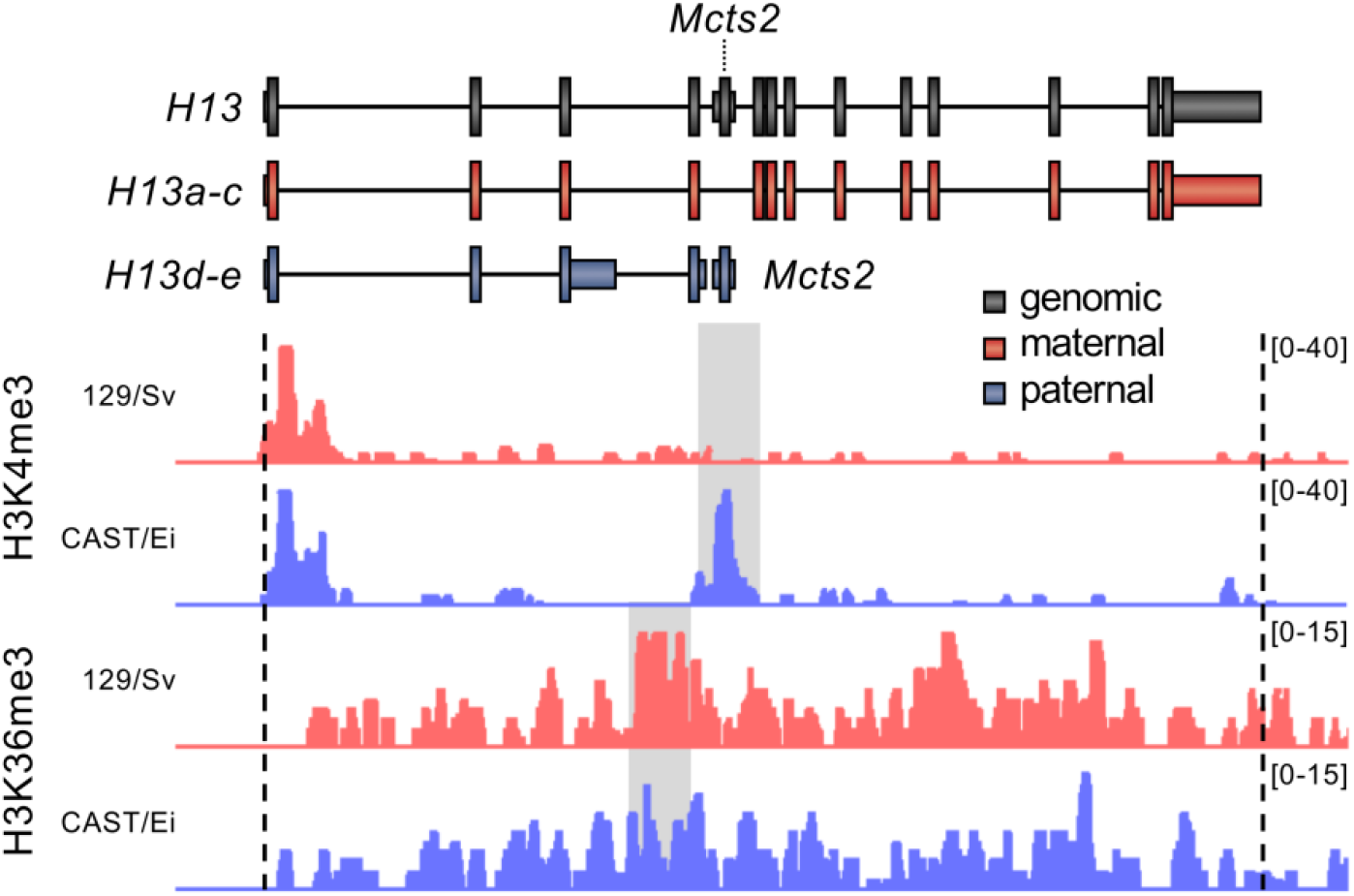
ChIP-seq profiles for H3K4me3 and H3K36me3 from hybrid immortalised mouse fibroblasts mapped to the maternal (129/Sv) or paternal (CAST/Ei) allele of *Mcts2/H13*. Genomic (black) and packed maternal (red) and paternal (blue) tracks are shown at the top. Coverage values have been normalised by input and are indicated on the y-axis. Allele-specific chromatin mark enrichments are highlighted in grey.

**Figure 4.**
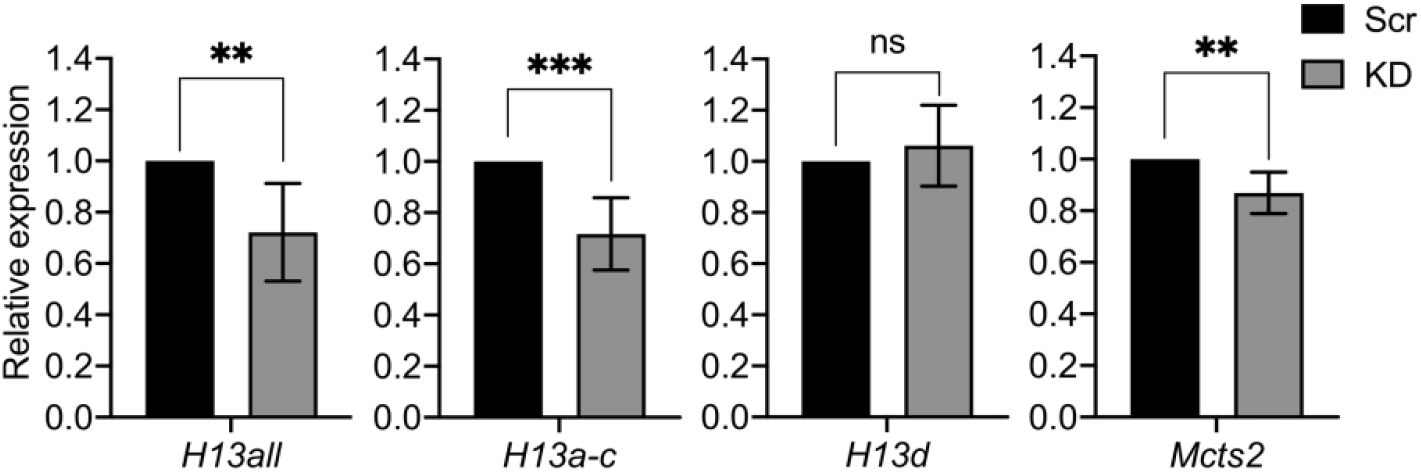
mRNA levels of target transcripts assessed by RT-qPCR 48 hours post transfection. All data are normalised to Ct values for *Actb*. Data are given as mean 2_−ΔΔCt_ values α SEM of three independent experiments. ns, p>0.05; **, p=0.01; ***, p<0.001 compared with Scrambled group by unpaired *t*-test. Scr, scrambled control; KD, knockdown.

### iCGI activity influences host gene transcription and IPA

The *Mcts2/H13* locus provides a model for studying APA and the role of iCGIs in alternative transcript formation. To understand the involvement of iCGI expression in transcriptome diversity more broadly, we devised a strategy to estimate the number of loci in the genome where iCGIs reside in host gene bodies and are associated with tissue-specific gene expression patterns. Tissue-specific variation in transcription at iCGIs was determined using ENCODE RNA-seq data for 30 mouse tissues and developmental stages (50). 4033 host genes were identified genomewide that harbour iCGIs located at least 1 kb from their TSS and 500 bp from the start of their last exon. A negative control dataset was generated consisting of 1079 well-defined loci with either a corresponding protein entry in PDB or validation by RefSeq that neither harbour an iCGI nor overlap with other genes. Then, an artificial iCGI was simulated at each of the negative control loci, with its position and size randomly drawn from the normalised position and size distributions of the actual iCGIs. For each iCGI/host gene pair identified in the genome, RNA-seq reads mapping *upstream*, *across* and at the *iCGI* were counted (Figure 5A). Additionally, the ratio of reads mapping *upstream* and *across* the iCGI was calculated for each gene harbouring an iCGI and used to determine the proportion of host gene transcripts terminating upstream of the iCGI or traversing the iCGI. Pearson correlation coefficients (ρ) were calculated between *upstream:across* ratios and the number of RNA-seq reads mapping to the iCGIs themselves.

**Figure 5.**
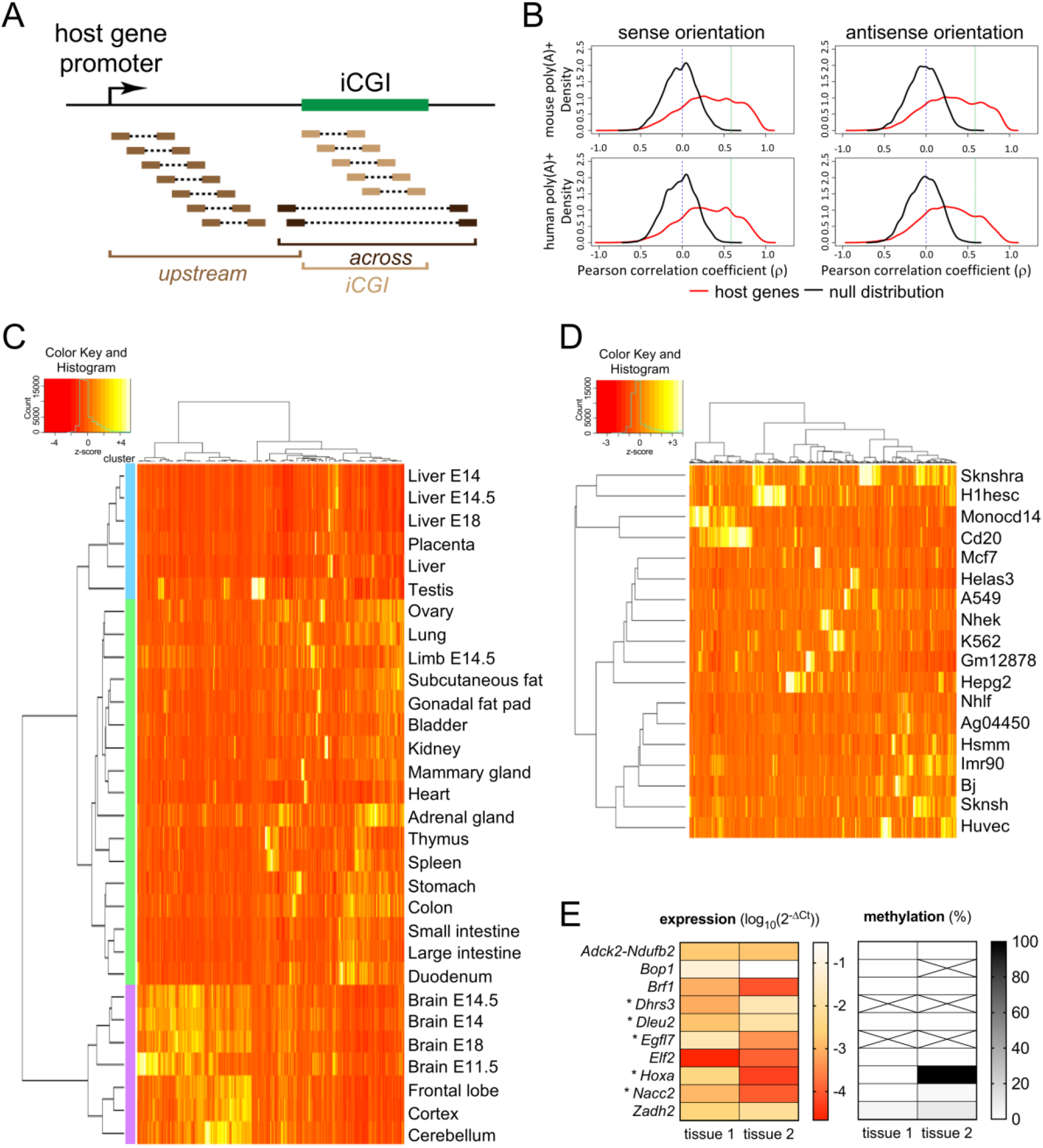
(A) Schematic representation of data collection. RNA-seq reads at iCGI/host gene pairs were divided into three groups according to the region they were mapped to: *upstream*, *across* or at the *iCGI*. (B) Pearson correlation coefficients (ρ) between transcription from the *iCGI* and transcription *upstream:across* the iCGI. ρ values were calculated in both sense (left) and antisense (right) orientations with respect to the host gene across 30 mouse tissues (upper) and 18 human cell lines (lower) using RNA-seq data from polyadenylated [poly(A)+] transcripts. A vertical blue dashed line is at ρ=0. A strict cut-off is represented by a vertical green dashed line at ρ=0.59, equal to the maximum ρ value observed in the null distribution. (C) RNA-seq heatmap illustrating tissue- and developmental stage-specific transcriptional activity of murine iCGIs within host genes with ρ>0.59. Values are given as column-wise standard-normalised fragments per kilobase of transcript per million mapped reads (z-score). Tissues are from adult mice, unless specified. (D) RNA-seq heatmap illustrating cell type-specific transcriptional activity of human iCGIs within host genes with ρ>0.59. Values are given as column-wise standard-normalised fragments per kilobase of transcript per million mapped reads (z-score). (E) Ten iCGIs from C were selected and labelled with the name of their host gene. Transcription from the iCGIs was measured by RT-qPCR in two tissues (left). All data are normalised to Ct values for *Actb*. Data are given as log_10_ of mean 2_−ΔCt_ values of three independent experiments. *, expression is consistent with RNA-seq data in C. DNA methylation was measured by sequencing of bisulfite converted genomic DNA and is given as percentage values (right). Crossed out cells indicate that DNA methylation could not be determined.

The distribution of ρ values of host genes with iCGIs was found to differ substantially from that of the negative control dataset (null distribution) (Figure 5B, *upper*). The null distributions are centred around zero and have a moderate range, indicating that this approach has a sufficient level of specificity and that extreme ρ values are unlikely to occur by chance. The distributions originating from iCGI/host gene pairs are skewed to the right (Figure 5B), showing that increased iCGI transcription positively correlates with increased *upstream:across* ratios. In order to select a subset of candidate loci for further investigation, a strict cut-off value of 0.59, equal to the maximum ρ value observed in the negative data set, was imposed: 1722 (21%) iCGI/host gene pairs scored above this threshold. The sensitivity of this approach was validated by the detection of the imprinted pair *Nap1l5/Herc3* (ρ=0.94) among the significant loci. The promoter of *Nap1l5* is an iCGI that, when transcriptionally active, leads to *Herc3* IPA (23). The same analysis was conducted using ENCODE RNA-seq data sets from 18 human cell lines (50) and resulted in similar findings (Figure 5B, *lower* and Supplementary Figure S5). Transcription from a large number of iCGIs positively correlates with *upstream:across* host gene transcription ratios.

Although the model *Mcts2/H13* locus is subject to genomic imprinting, we have shown through this analysis that the majority of the identified iCGI/host gene pairs (over 4000) are not imprinted, suggesting that iCGI influence on host gene polyadenylation is widespread and conserved across two mammalian species.

### iCGIs show DNA methylation-independent spatiotemporal transcriptional activity

The transcriptional activity at iCGIs from iCGI/host gene pairs with ρ>0.59 was compared across all tissues, developmental stages and cell lines. Both mouse and human datasets showed striking tissue-specific iCGI activity with iCGIs often being highly expressed in a single tissue or cell line (Figure 5C and 5D). Hierarchical clustering grouped mouse tissues and developmental stages from the same organ or system within the same cluster (Figure 5C). The hierarchical clustering was reproducible when using only iCGI/host gene pairs with iCGIs fully mapping within one intron of their host genes, excluding iCGIs partially or fully overlapping with host gene exons (Supplementary Figure S6A). This is important since it is not possible to discriminate between same-strand RNA-seq reads from host genes and iCGIs when the latter fully or partially overlap with host gene exons. Additionally, when intronic iCGIs were subjected to hierarchical clustering according to their expression level, a large number of iCGIs that had high expression in brain tissues and developmental stages grouped together (Supplementary Figure S6A). GO term analysis using host genes harbouring these iCGIs revealed enrichment of brain-specific biological processes (Supplementary Figure S6B), confirming that these host genes are involved in relevant cell specification processes.

In order to establish a role for DNA methylation in the regulation of iCGIs transcriptional activity in mouse, ten candidate iCGI/host gene pairs were selected for further analysis. For each iCGI, two tissues were assayed that exhibited the highest and lowest measured expression levels based on RNA-seq data. For five of the iCGIs, the gene expression measured by RT-qPCR recapitulated the gene expression patterns detected in the RNA-seq datasets (Figure 5E, *left* and Supplementary Figure S7) which is consistent with expectations. The other five iCGIs did not match the RNA-seq expression patterns using RT-qPCR. However, these RT-qPCR assays were undertaken in tissues that were as close as possible to the descriptions in the ENCODE data repository but were not exact matches to tissues used by the ENCODE consortium. Therefore, there were differences in the precise age of collected embryos in developmental samples and differences in the dissected sections of particular tissues, possibly accounting for the discordance. In some cases, strain differences may also have led to discrepant findings. We subsequently assessed the methylation status of these candidate loci by PCR amplification of bisulfite-converted genomic DNA. At one of the concordant iCGI/host gene pairs, namely *Hoxa*, DNA methylation correlated with gene expression (Figure 5E, *right*). For the other four, no correlation between iCGI expression and DNA methylation was detected (Figure 5E, *right*) suggesting that DNA methylation is not the major regulatory factor involved.

### Active intronic iCGIs present promoter-like chromatin

To determine whether these iCGIs are independent transcriptional units or a by-product of host gene transcription, ENCODE H3K36me3 and H3K4me3 ChIP-seq datasets from mouse E14.5 brain (50) were interrogated. When in promoter regions, H3K36me3 negatively affects transcription (60), whereas H3K4me3 is considered a typical promoter mark and it can be found at both transcriptionally active (61) and poised promoters (62). In order to identify appropriate subsets for comparison, hierarchical clustering was applied to host gene expression upstream of the iCGI and the expression of the iCGI itself. This provided a means to select transcriptionally active host genes with very active or less active iCGIs and exclude iCGIs within inactive host genes from further analysis (Figure 6A). Furthermore, before plotting average gene body H3K36me3 and H3K4me3 profiles, only host genes with iCGIs fully contained within introns were selected since H3K36me3 is enriched at expressed exons (12). This approach revealed a sharp decrease in H3K36me3 around highly active iCGIs and a more subtle depletion around iCGIs expressed at low levels (Figure 6B, *upper*). H3K36me3 gradually accumulates downstream of highly expressed iCGIs before decreasing again upstream of host gene transcription termination sites (TTSs), resembling the deposition of H3K36me3 along the body of actively transcribing genes (Figure 6B, *upper left*). H3K4me3 was enriched at all intronic iCGIs with a bias towards highly expressed iCGIs (Figure 6B, *lower*). Taken together, these findings suggest that the iCGIs tested may indeed be discrete tissue-specific promoters and that histone modifications could be important to regulate their expression.

**Figure 6.**
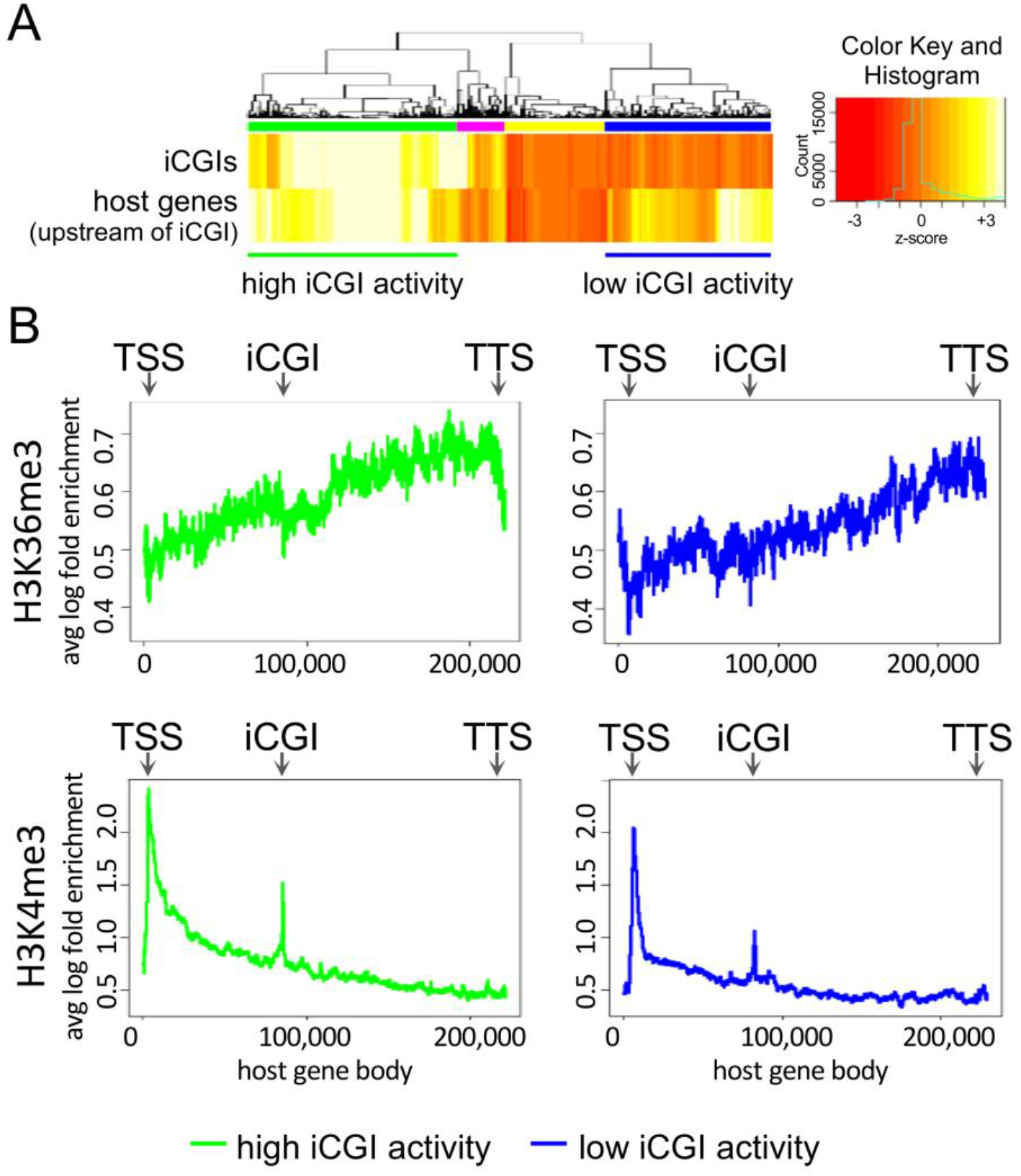
(A) RNA-seq heatmap from E14.5 mouse brain illustrating the transcriptional activity of iCGI/host gene pairs showing a ρ>0.59 between transcription from the *iCGI* and transcription *upstream:across* of the iCGI (see main text and Figure 5A). Highly expressed iCGIs within active or inactive host genes are highlighted with green or pink bars, respectively. Moderately expressed iCGIs within active or inactive host genes are highlighted with blue or yellow bars, respectively. Values are given as column-wise standard-normalised fragments per kilobase of transcript per million mapped reads (z-score). (B) Average H3K36me3 (upper) and H3K4me3 (lower) ChIP-seq profiles for iCGI/host gene pairs from A in which the iCGIs are fully intronic. Reads are mapped to the body of active host genes harbouring intronic iCGIs that are transcribed at high (left) or low (right) levels. TSS, transcription start site; TTS, transcription termination site.

## DISCUSSION

Epigenetic modifications are associated with transcription and pre-mRNA processing. In some cases, the deposition of one mark is dependent upon the presence or absence of another. The same marks can differentially influence transcription depending on genomic context. For instance, DNA hypermethylation at CGI-associated promoters negatively impacts transcription (34) but facilitates it when present within gene bodies (32). One post-translational histone modification has been extensively studied in the context of pre-mRNA processing, namely H3K36me3 (14, 16, 28, 29, 54). This histone mark is deposited co-transcriptionally along the body of actively transcribing genes by the RNA Pol II-interacting histone methyltransferase SETD2 (51). Intriguingly, human kidney tumours with *SETD2* mutations are characterised by increased intron retention and altered transcription termination site usage (28, 29).

We investigated the influence of H3K36me3 on intron retention and IPA in NIH/3T3 cells using siRNAs coupled with transcriptome analysis and locus-specific gene expression assays. We show that depletion of H3K36me3 leads to significant deregulation of 4266 transcripts and 136 AS events. Changes in intron retention are virtually absent between *Setd2* KD and WT samples, indicating that H3K36me3 is not a master regulator of this type of pre-mRNA processing. Locus-specific experiments on the imprinted iCGI/host gene pair *Mcts2/H13* are consistent with our genomewide conclusions since we did not detect changes in *H13d* transcript abundance, a proxy for intron retention, upon H3K36me3 depletion. Our findings confirm the involvement of H3K36me3 in AS but differ from previous studies in human kidney tumours that associate *SETD2* mutations with increased intron retention. This could be partially explained by our use of polyadenylated RNA rather than total RNA for the preparation of RNA-seq libraries. If H3K36me3 depletion does indeed lead to increased intron retention in the nucleus, there must be efficient downstream checkpoints that prevent the export of those transcripts into the cytoplasm.

Since paternal *H13* transcripts are characterised by intron retention and IPA upstream of an actively transcribing iCGI, we interrogated the impact of transcription from iCGIs on host gene pre-mRNA processing more generally. We bioinformatically analyse 30 mouse tissues and developmental stages using RNA-seq datasets from the ENCODE project and identify 4033 iCGI/host gene pairs. For one in five pairs, iCGI activity is tissue- or developmental stage-specific and the level of host gene transcription upstream of the iCGI positively correlates with the level of iCGI transcription. We repeated the same analysis using ENCODE RNA-seq data from 18 human cell lines and find the same results, indicating that these observations are reproducible across two mammalian species. Additionally, we demonstrate that this effect is largely independent of DNA methylation for a small subset of iCGI/host gene pairs. Finally, we provide evidence that these iCGIs may be discrete tissue-specific promoters since they are enriched for H3K4me3 and depleted of H3K36me3, a chromatin profile typically associated with active gene promoters (63).

Previous studies have shown that iCGIs become methylated and silenced when host gene transcription traverses them and triggers the recruitment of the *de novo* DNA methylation machinery (33). However, the methylation status of these iCGIs is dependent upon their intrinsic ability to initiate transcription, i.e. strong iCGIs can escape host gene transcription-mediated silencing (33). These iCGIs retain an unmethylated state, are enriched in H3K4me3 and lack H3K36me3 (33). Our work consolidates these findings and provides an additional layer of complexity to the cross talk between host gene promoters and iCGIs. We show for the first time that transcription from a large number of iCGIs interferes with host genes transcription in mouse and human, possibly providing a novel mechanism for spatiotemporal diversification of both transcriptome and proteome. These findings raise the question of precisely how iCGIs influence pre-mRNA processing of host gene transcripts. We speculate that iCGI activity may stimulate IPA at the expense of 3’UTR-APA in host genes with multiple polyadenylation sites. Isoforms resulting from IPA are typically as robustly expressed as full-length transcripts and therefore likely represent functional mRNAs rather than transcriptional noise (11). This is particularly relevant since IPA is important for regulating transcript diversity during differentiation and development in both physiological and pathological conditions (11). Using differentiation models, it will be important to examine the extent of the role of iCGIs in tissue- and developmental stage-specific gene expression and to understand the mechanisms involved.

## AVAILABILITY

The splice donor site score calculation tool is freely available at http://rulai.cshl.edu/new_alt_exon_db2/HTML/score.html.

## ACCESSION NUMBERS

*Setd2* knockdown RNA-seq data have been deposited to GEO under accession number GSE147077.

## ACKNOWLEDGEMENTS

The authors acknowledge the support from the Department of Health via the National Institute for Health Research, Biomedical Research Centre at Guy’s and St Thomas’ NHS Foundation Trust in partnership with King’s College London for access to next generation sequencing and use of RT-qPCR equipment. We also thank the King’s College London genomics core facility for access to Sanger sequencing equipment.

