## Supplementary Data for "Transcription of intragenic CpG islands influences spatiotemporal host gene pre-mRNA processing"

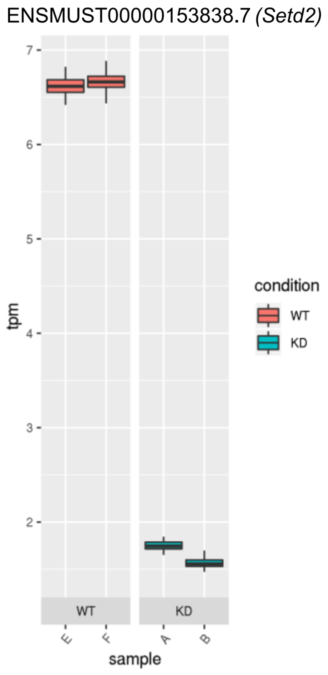


**Supplementary Figure S1.** Expression of *Setd2* (ENSMUST00000153838.7) in wild type (WT) and knockdown (KD) samples. Data are given as normalised RNA-seq transcript counts (tpm).

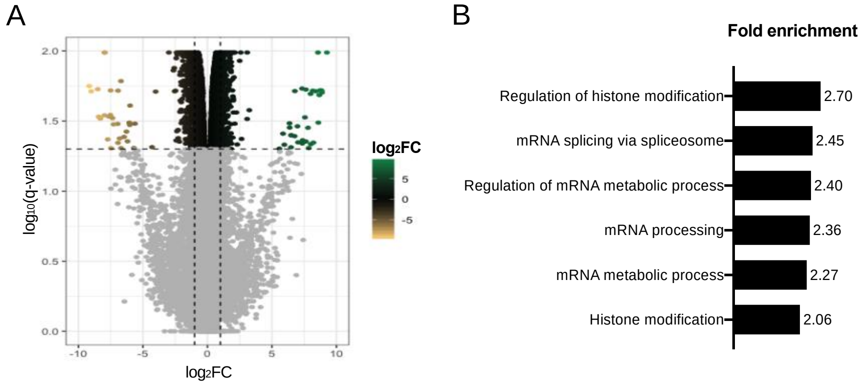


**Supplementary Figure S2.** (A) RNA-seq volcano plot illustrating differentially expressed transcripts from knockdown samples. Two vertical dashed lines are at $\log_2 FC = \pm 1$ and one horizontal dashed line is at q-value = 0.05. (B) Upregulated biological processes determined by GO analysis (PANTHER). See Supplementary Table S5 for a complete list of GO ID terms.

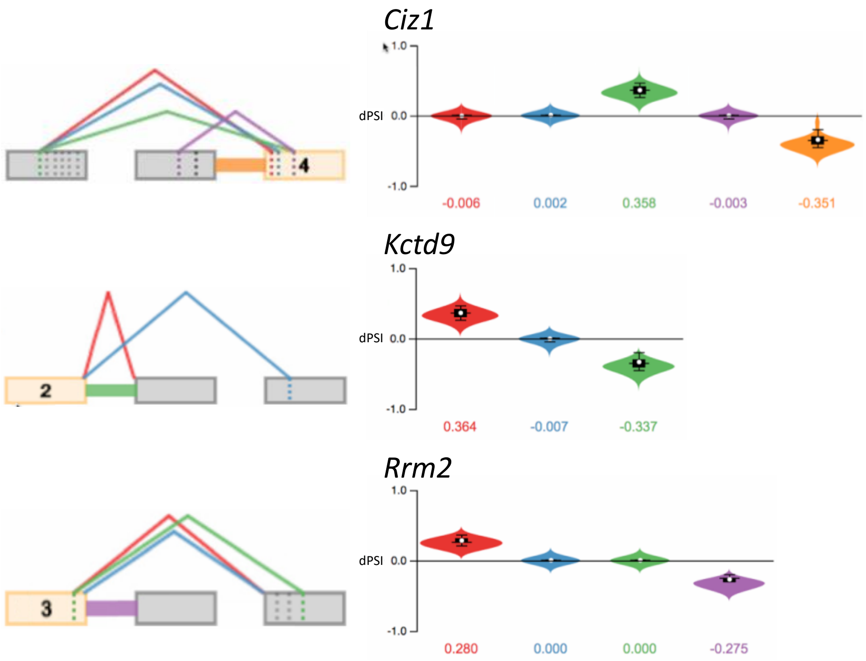


**Supplementary Figure S3.** Local splice variants (LSVs) plots generated by MAJIQ/Voila. For each gene, all significant LSVs are shown on the left and their associated relative usage values (dPSI) on the right in the same colour. A positive dPSI value indicates that the LSV is used more in *Setd2* knockdown samples and vice versa.

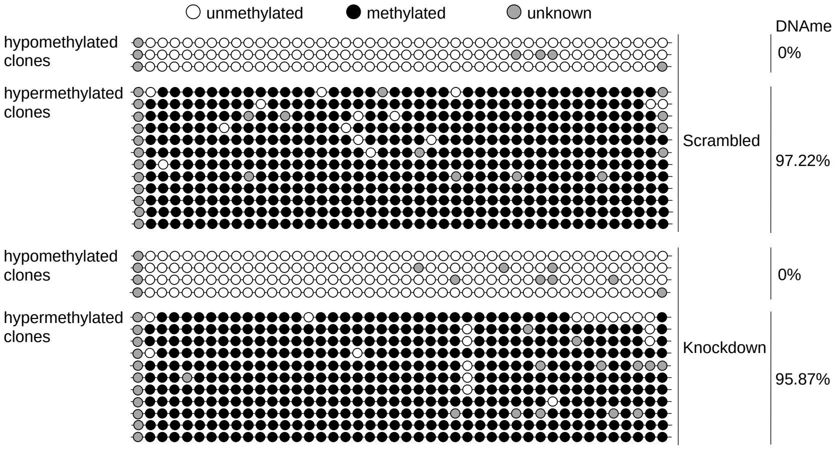


**Supplementary Figure S4.** *Mcts2/H13* iCGI methylation upon *Setd2* knockdown. Horizontal lines represent individual strands of DNA and circles represent cytosine residues in a CpG context. DNA methylation (DNAme) percentage values are also shown. White circles, unmethylated CpGs; black circles, methylated CpGs; grey circles, the methylation status of the CpG is unknown.

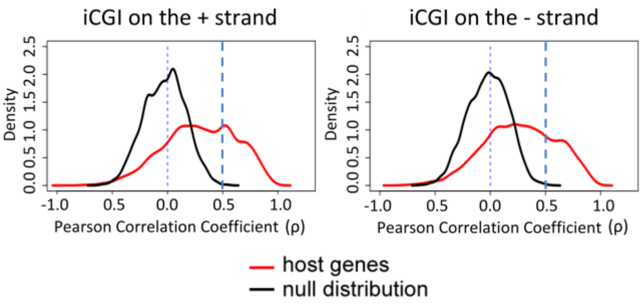


**Supplementary Figure S5.** Pearson correlation coefficients (ρ) between transcription from the *iCGI* and transcription *upstream:across* the iCGI (see main text and Figure 5A). ρ values were calculated in both sense (left) and antisense (right) orientations with respect to the host gene across 18 human cell lines using RNA-seq data from non-polyadenylated transcripts. A vertical blue dashed line is at ρ=0. A strict cut-off is represented by a second vertical blue dashed line at ρ=0.59, equal to the maximum ρ value observed in the null distribution.

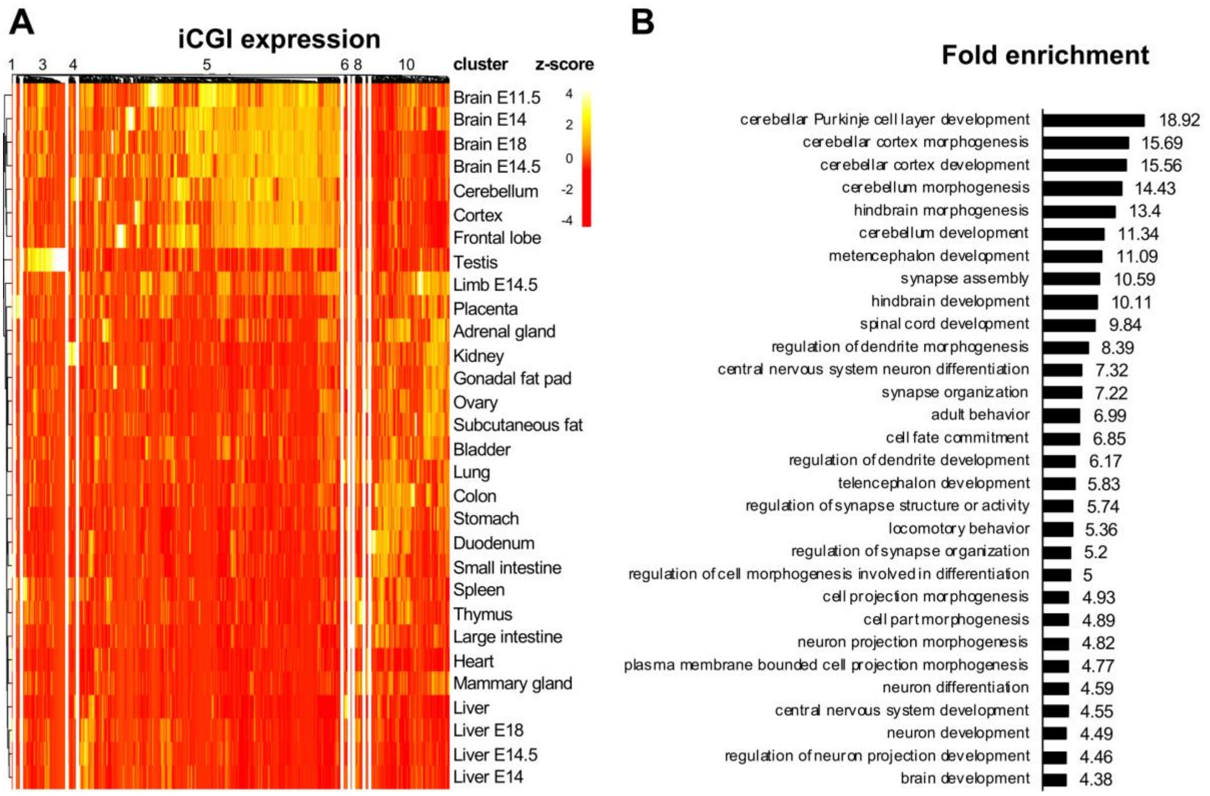


**Supplementary Figure S6.** (A) RNA-seq heatmap illustrating tissue- and developmental stage-specific transcriptional activity of murine intronic iCGIs within host genes with ρ>0.59 (see Figure 5B). Values are given as column-wise standard-normalised fragments per kilobase of transcript per million mapped reads (z-score). Tissues are from adult mice, unless specified. (B) Upregulated biological processes determined by GO analysis (PANTHER) using host genes harbouring the intronic iCGIs grouped in cluster 5 (see A). See Supplementary Table S7 for a complete list of GO ID terms.

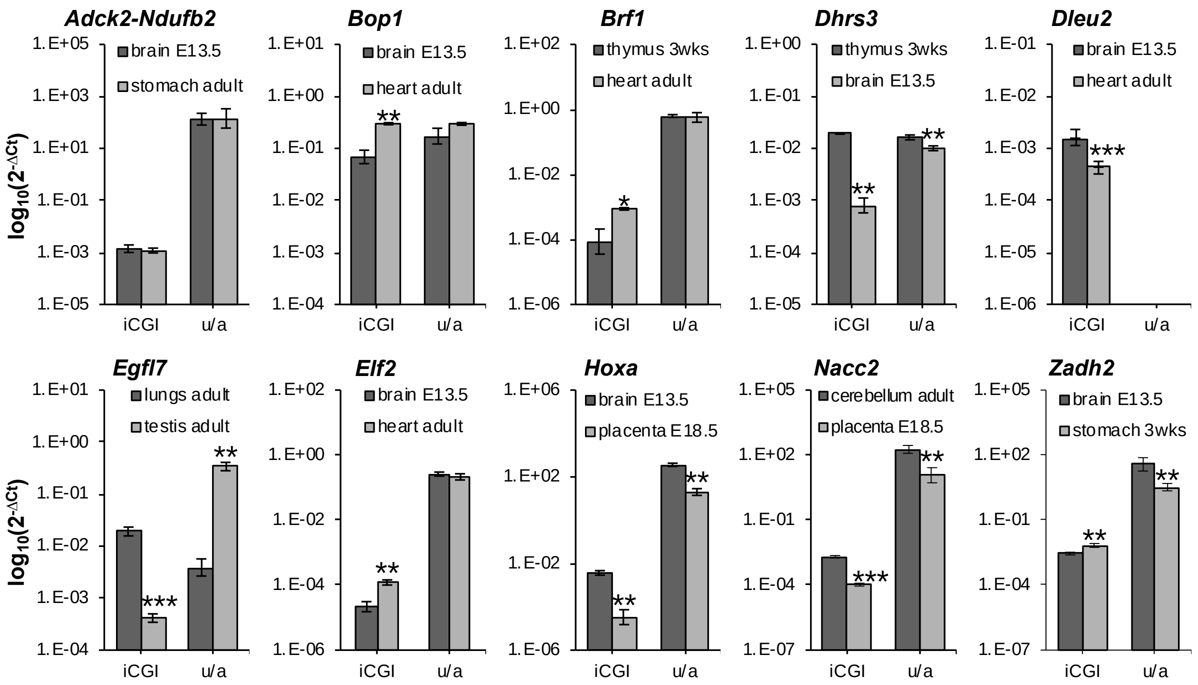


**Supplementary Figure S7.** mRNA levels of target iCGIs and relative host genes *upstream:across* ratios (u/a) assessed by RT-qPCR. All data are normalised to Ct values for *Actb*. Data are given as $\log_{10}$ of mean $2^{-\Delta Ct}$ values ± 95% confidence interval of three independent experiments. *, p<0.05; **, p=0.01; ***, p<0.001 compared with the other tissue by unpaired *t*-test.

**Supplementary Table S1.** Antibodies

| Target | Manufacturer | Cat. No | Dilution |
|---|---|---|---|
| ACTB | Applied Biosystems | 8457 | 1:1000 |
| H3 | Applied Biosystems | ab1791 | 1:1500 |
| H3K36me3 | Applied Biosystems | ab9050 | 1:500 |
| IgG | Applied Biosystems | 7074 | 1:10000 |

**Supplementary Table S2.** TaqMan probes

| Target | Manufacturer | Assay ID |
|---|---|---|
| *Actb* | Applied Biosystems | Mm00607939_s1 |
| *H13* | Applied Biosystems | Mm00468785_m1 |
| *H13a-c* | Applied Biosystems | Mm01241449_m1 |
| *H13d* | Applied Biosystems | Custom |
| *Mcts2* | Applied Biosystems | Mm00481540_s1 |
| *Setd2* | Applied Biosystems | Mm01250225_m1 |

**Supplementary Table S3.** Primers

| Name | Sequence | Experiment |
|---|---|---|
| iMcts2F1 | TTTTTAAGTATTAGAATATTGGGGGATT | 2nd round nested PCR |
| iMcts2R1 | AACATAATCTTAATAAAAAAACACC | 2nd round nested PCR |
| oMcts2F1 | TTTTTGGTTGTTAAGTATATTTTTGT | 1st round nested PCR |
| oMcts2R1 | TACAATTAAACACACTTTCCTTCTC | 1st round nested PCR |
| SP6 | ATTTAGGTGACACTATAG | Bisulfite sequencing |
| T7 | TAATACGACTCACTATAGGG | Bisulfite sequencing |

**Supplementary Table S4.** *H13* splice donor sites scores

| Exon | Score | Exon | Score |
|---|---|---|---|
| 1 | 8.1 | 7 | 9.2 |
| 2 | 9.7 | 8 | 8.8 |
| 3 | 4.4 | 9 | 10.4 |
| 4 | 5.9 | 10 | 10.1 |
| 5 | 12.6 | 11 | 11 |
| 6 | 8.8 | 12 | 8.1 |
